# Flavonol rhamnosylation indirectly modifies the cell wall defects of *RHAMNOSE BIOSYNTHESIS1* mutants by altering rhamnose flux

**DOI:** 10.1101/175133

**Authors:** Adam M. Saffer, Vivian F. Irish

## Abstract

Rhamnose is required in *Arabidopsis thaliana* for synthesizing pectic polysaccharides and glycosylating flavonols. *RHAMNOSE BIOSYNTHESIS1 (RHM1)* encodes a UDP-L-rhamnose synthase, and *rhm1* mutants exhibit many developmental defects, including short root hairs, hyponastic cotyledons, and left-handed helically twisted petals and roots. It has been proposed that the hyponastic cotyledons observed in *rhm1* mutants are a consequence of abnormal flavonol glycosylation, while the root hair defect is flavonol-independent. We have recently shown that the helical twisting of *rhm1* petals results from decreased levels of rhamnose-containing cell wall polymers. In this work, we find that flavonols indirectly modify the *rhm1* helical petal phenotype by altering rhamnose flux to the cell wall. Given this finding, we further investigated the relationship between flavonols and the cell wall in *rhm1* cotyledons. We show that decreased flavonol rhamnosylation is not responsible for the cotyledon phenotype of *rhm1* mutants. Instead, flavonols provide a large reservoir of rhamnose, and blocking flavonol synthesis or rhamnosylation can suppress *rhm1* defects by diverting rhamnose to the synthesis of cell wall polysaccharides. Therefore, rhamnose is required in the cell wall for normal expansion of cotyledon epidermal cells. Our findings suggest a broad role for rhamnose-containing cell wall polysaccharides in the morphogenesis of epidermal cells.

## Introduction

Carbohydrates are important for all aspects of plant biology. Many different polysaccharides are critical components of the plant cell wall (Cosgrove, 2005). Mono-and oligosaccharides can also act as signaling molecules (Gibson, 2005), and can modify the properties of other molecules such as proteins and metabolites by glycosylation (Bowles et al., 2005; Strasser, 2016).

Plant cell walls are important for defense (Bethke et al., 2016), cell-cell adhesion (Daher and Braybrook, 2015), structural support, and the control of cell morphology (Cosgrove, 2005). The cell wall is largely composed of polysaccharides, including cellulose, hemicelluloses and pectins. Cellulose is a polymer of (1,4)-linked β-D-glucans, which are bundled together as microfibrils that provide mechanical support and control the direction of cell expansion. Cellulose microfibrils are crosslinked by hemicelluloses, such as xyloglucans and xylans (Cosgrove, 2005). Cellulose and hemicelluloses are embedded in a gel-like matrix of pectin. Pectin is a complex mix of different polysaccharides, including homogalacturonan (HG), rhamnogalacturonan-I (RG-I), and rhamnogalacturonan-II (RG-II), all of which are likely crosslinked by a mix of covalent and non-covalent interactions (Mohnen, 2008). The most abundant pectic polysaccharide is HG, a polymer of (1,4)-linked α-D-galacturonic acid that can be modified by methylesterification on C6 (Mohnen, 2008). RG-II is a complex and conserved polysaccharide consisting of a short galacturonic acid backbone decorated by four sidechains, incorporating twelve different monosaccharides, including several rhamnose residues in the sidechains (O’Neill et al., 2004). RG-II is present in the cell wall as a borate crosslinked dimer that is essential for plant growth (O’Neill et al., 2001). RG-I is the second most abundant pectic polysaccharide, and consists of a repeating disaccharide (1,2)-α-L-rhamnose-(1,4)-α-D-galacturonic acid backbone with many of the rhamnose residues substituted with galactan, arabinan, and arabinogalactan sidechains (Mohnen, 2008).

Flavonoids, including the flavonols kaempferol, quercetin, and isorhamnetin, are a major class of secondary metabolites that are frequently glycosylated. Flavonoids provide UV protection (Li et al., 1993), regulate auxin signaling (Brown et al., 2001; Buer and Muday, 2004; Peer et al., 2004; Yin et al., 2014), and in some species are required for male fertility (Burbulis et al., 1996; Mo et al., 1992). In Arabidopsis, flavonols can be glycosylated at the C3 and C7 positions, primarily with rhamnose and glucose (Kachlicki et al., 2008; Kerhoas et al., 2006; Ringli et al., 2008; Routaboul et al., 2006; Yin et al., 2012; Yonekura-Sakakibara et al., 2008). Flavonol glycosylation can be tissue-specific; for example, flowers have particularly high levels of flavonols and there are many flower-specific flavonol glycosides (Yonekura-Sakakibara et al., 2008; Yonekura-Sakakibara et al., 2007). In most tissues essentially all flavonols are glycosylated, and if flavonol glycosylation is genetically blocked plants can downregulate flavonol synthesis to prevent flavonol aglycones from accumulating (Yin et al., 2012). Specific flavonol glycosides might function as auxin transport inhibitors or in herbivore defense (Onkokesung et al., 2014; Yin et al., 2014).

Rhamnose is critical for synthesis of the pectic polysaccharides RG-I and RG-II and for flavonol glycosylation. UDP-L-rhamnose synthase enzymes convert UDP-D-glucose to UDP-L-rhamnose, which is the rhamnosyl donor used by glycosyltransferases (Jones et al., 2003; Yonekura-Sakakibara et al., 2007). UDP-L-rhamnose synthases are encoded by three highly similar genes in Arabidopsis, *RHAMNOSE BIOSYNTHESIS1/REPRESSOR OF LRX1 (RHM1/ROL1)*, *RHM2/MUM4*, and *RHM3* (Oka et al., 2007; Reiter and Vanzin, 2001). The *RHM2/MUM4* gene product primarily synthesizes UDP-L-rhamnose for RG-I in seed mucilage (Usadel et al., 2004; Western et al., 2004), while the developmental role of *RHM3* has not yet been determined.

*RHM1/ROL1* (hereafter referred to as *RHM1*) is required for morphogenesis and expansion of multiple epidermal cell types, including root hairs, cotyledon pavement cells, and conical petal epidermal cells (Diet et al., 2006; Ringli et al., 2008; Saffer et al., 2017). *rhm1* mutant adaxial cotyledon pavement cells have cell expansion defects and fail to form their characteristic lobes, leading to hyponastic cotyledons (Ringli et al., 2008). *RHM1* is required for the synthesis of rhamnose-containing cell wall polymers that prevent left-handed helical twisting of root and petal cells (Saffer et al., 2017). *rhm1* mutants decrease the overall abundance of cell wall rhamnose in petals, reduce levels of the pectic polysaccharides RG-I and RG-II in roots, and accumulate non-rhamnosylated flavonols in multiple tissues (Diet et al., 2006; Ringli et al., 2008; Saffer et al., 2017). Blocking flavonol synthesis suppresses some *rhm1* defects including hyponastic cotyledons, which has led to the hypothesis that decreased flavonol rhamnosylation is responsible for the *rhm1* cotyledon defects (Ringli et al., 2008). By contrast, other *rhm1* defects such as abnormal root hairs are unaffected by loss of flavonols, and so these phenotypes were proposed to be a consequence of altered cell wall composition (Ringli et al., 2008).

Because rhamnose is a component of both pectic polysaccharides and flavonols, impairing the synthesis or modification of either class of molecules has the potential to alter rhamnose flux and indirectly affect the other class. Here, we show that decreased flavonol rhamnosylation does not cause the helical petal or hyponastic cotyledon phenotypes of *rhm1* mutants. Instead, decreased levels of pectic polysaccharides likely cause the *rhm1*-associated abnormal cell morphology defects, and blocking flavonol synthesis or flavonol rhamnosylation can suppress those defects by diverting rhamnose from flavonols to the cell wall.

## Results

### *rhm1* decreases flavonol rhamnosylation in flowers

*rhm1* mutants have helically twisted petals and cell expansion defects in adaxial petal epidermal cells (Saffer et al., 2017). To investigate the role of *RHM1* in flavonol glycosylation in flowers, we used HPLC-MS to analyze the abundance of flavonol glycosides in flowers of L*er* wild-type and the *rhm1-3* mutant. Peaks representing 17 different flavonol glycosides were identified (Figure 1A). Most flavonol glycosides were present in both genotypes, although some were specific to either wild-type or *rhm1-3* flowers. Based on mass, retention time, and comparison to a previous description of floral flavonol glycosides (Yonekura-Sakakibara et al., 2008), we were able to identify the specific flavonol and conjugated sugars for 13 of these flavonol glycosides (Figure 1B). The *transparent testa 4* (*tt4*) mutant is defective in chalcone synthase and consequently lacks all flavonoids, including flavonols (Burbulis et al., 1996). All compounds that we had identified as flavonols were absent in extracts from *rhm1-3 tt4* double mutant flowers, confirming that they were flavonoids (Figure 1A). Several peaks were still present in the *rhm1-3 tt4* double mutant, but in each case the peak in *rhm1-3 tt4* represented a compound with different mass and UV absorption spectra compared to *rhm1-3* or the wild-type. For example, quercetin-3-O-glucoside (Q-3G) and sinnapoyl malate co-eluted, and for wild-type and *rhm1-3* flowers the peak consisted primarily of Q-3G. In the *rhm1-3 tt4* double mutant there was a substantial peak present that overlapped with the Q-3G peak, but the UV absorption spectrum was different than in the other genotypes, and the [M+H]+ mass of 341 was consistent with the peak primarily representing sinnapoyl malate.

**Figure 1.**
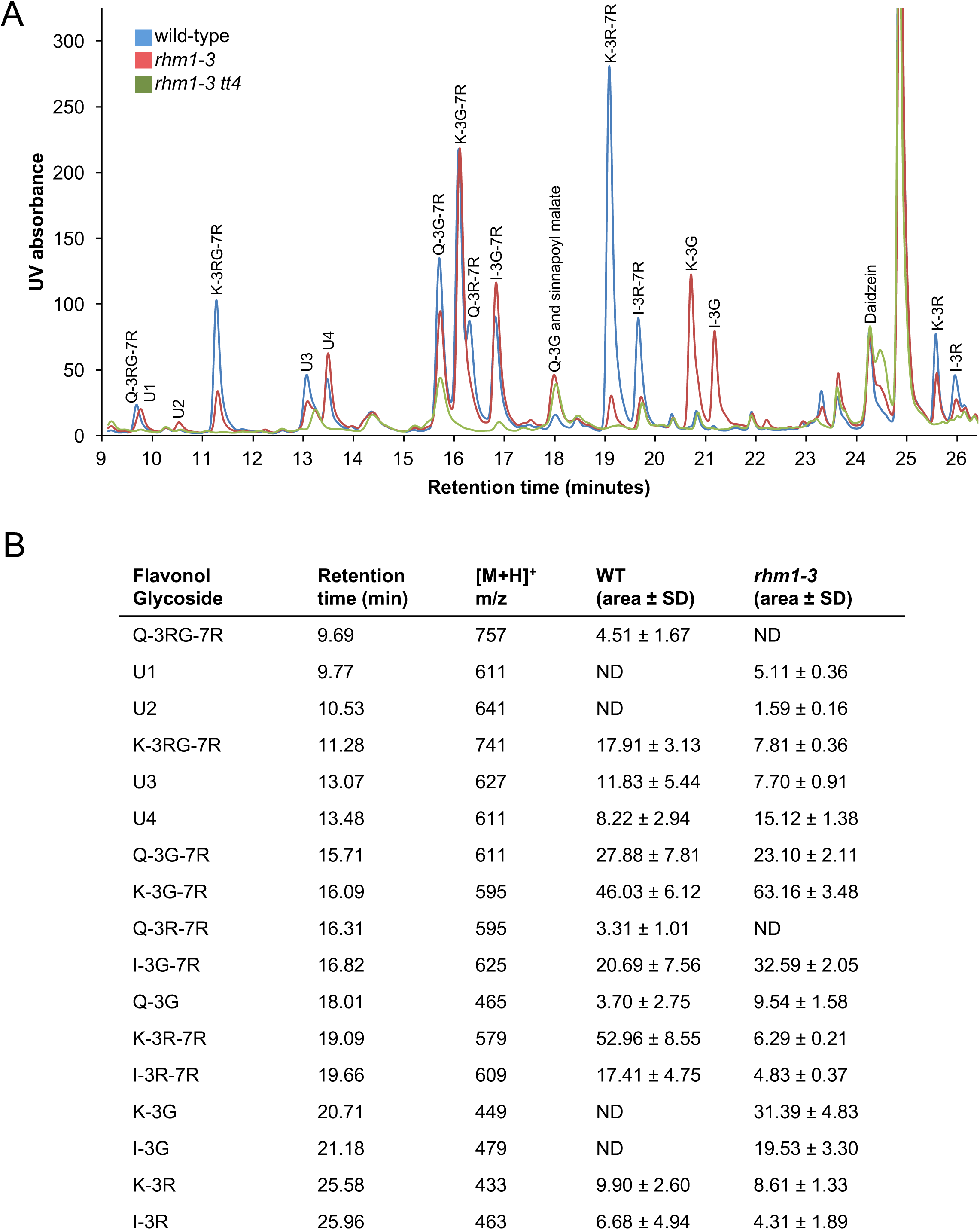
*rhm1-3* alters flavonol glycosylation in flowers. A) HPLC-MS analysis of methanol extracts from flowers. Flavonol abundance was determined by UV absorbance at 280 nm. L*er* wild-type*, rhm1-3*, and *rhm1-3 tt4* double mutant traces are each an average of three independent samples. Peaks that represent flavonols as determined by UV absorbance spectra and mass are labeled with the specific flavonol and the sugars conjugated to the 3 and 7 positions. U1-U4 represent peaks that appear likely to be flavonols based on UV absorbance spectra but for which the specific flavonol glycoside could not be unambiguously assigned. B) The peak areas in (A) were calculated to determine the relative abundance of each flavonol glycoside in wild-type and *rhm1-3* flowers. ND = not detected. Kaempferol (K), Quercetin (Q), Isorhamnetin (I), Glucose (G), Rhamnose (R).

In wild-type flowers, flavonols were glycosylated primarily with rhamnose and glucose, and the most abundant flavonol glycosides all included one or two rhamnose sugars, including kaempferol-3-O-rhamnoside-7-O-rhamnoside (K-3R-7R) and kaempferol-3-O-glucoside-7-O-rhamnoside (K-3G-7R) (Figure 1). Approximately half of identified flavonol glycosides in wild-type flowers were rhamnosylated at the C3 position, while 90% were rhamnosylated at the C7 position (Supplemental Table 1). The flavonol glycosylation profile of *rhm1-3* mutant flowers differed, with a substantial reduction in all flavonol di-rhamnosides, including K-3R-7R, quercetin-3-O-rhamnoside-7-O-rhamnoside (Q-3R-7R), and isorhamnetin-3-O-rhamnoside-7-O-rhamnoside (I-3R-7R). There was a corresponding increase in non-rhamnosylated flavonols in *rhm1-3*, including substantial amounts of kampferol-3-O-glucoside (K-3G) and isorhamnetin-3-O-glucoside (I-3G), which were undetectable in wild-type flowers. The levels of most single rhamnoside species, such as K-3G-7R, were similar between wild-type and *rhm1-3*. The effect of *rhm1-3* on flavonol glycosylation in flowers is similar to the effects previously reported for other *RHM1* mutations on flavonol glycosylation in flowers and seedlings (Ringli et al., 2008; Yonekura-Sakakibara et al., 2008). *rhm1-3* had a more severe effect on 3-O-rhamnosylation than 7-O-rhamnosylation, with a 72% decrease in the amount of 3-O-rhamnosylated flavonols and only a 28% decrease in the amount of 7-O-rhamnosylated flavonols (Supplemental Table 1). As a result, the ratio of 7-O-rhamnose to 3-O-rhamnose was increased from 1.69 in wild-type to 4.33 in *rhm1-3*. The *rhm1-3* mutation did not significantly alter overall levels of flavonols; the area of all peaks corresponding to putative flavonols was 231 ± 49 for wild-type and 241 ± 23 for *rhm1-3*.

### Loss of flavonols suppresses *rhm1* petal defects and diverts rhamnose to the cell wall

Blocking flavonol biosynthesis with a *tt4* mutant suppresses a subset of the cell morphology defects of *rhm1* mutant seedlings (Ringli et al., 2008). To test if flavonols contribute to the morphological defects seen in *rhm1* petals, we generated double mutant lines of *tt4* with *rhm1-3* or *rhm1-2* (also known as *rol1-2*). Wild type petals have a flat claw and blade (Fig 2A,B); in contrast, *rhm1-3* petal blades twist into a left-handed helix due to the left-handed twisting of their adaxial petal epidermal cells (Figure 2C; (Saffer et al., 2017)). *rhm1-2* has a more severe effect on petal development; most adaxial epidermal cells fail to expand causing a hyponastic petal phenotype and only occasional helical twisting (Figure 2D; (Saffer et al., 2017)). The *tt4* single mutant and *rhm1-3 tt4* and *rhm1-2 tt4* double mutants each had phenotypically normal petals with no helical twisting or hyponasty, indicating that loss of flavonols can restore normal development to *rhm1* petals (Figure 2E-G).

**Figure 2.**
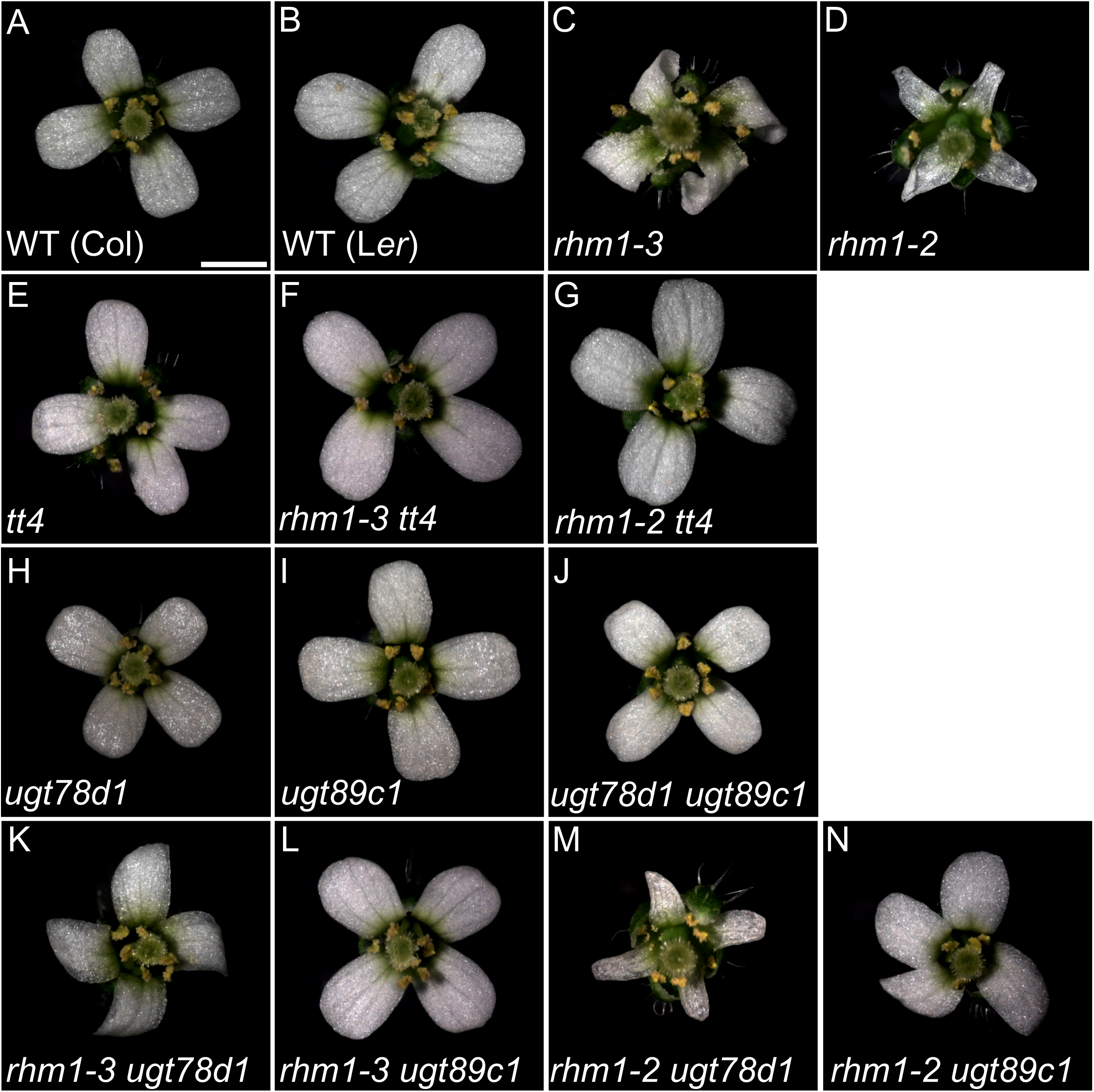
Blocking flavonol synthesis or rhamnosylation suppresses *rhm1* petal defects. Images of stage 14 flowers. (A) Col wild-type. (B) L*er* wild-type. (C) *rhm1-3* with left-handed twisted petals. (D) *rhm1-2* with hyponastic petals. (E) *tt4*. (F) *rhm1-3 tt4* double mutant. (G) *rhm1-2 tt4* double mutant. (H) *ugt78d1*. (I) *ugt89c1*. (J) *ugt78d1 ugt89c1* double mutant. (K) *rhm1-3 ugt78d1* double mutant. (L) *rhm1-3 ugt89c1* double mutant. (M) *rhm1-2 ugt78d1* double mutant. (N) *rhm1-2 ugt89c1* double mutant. Scale bar is 1 mm for all panels.

The hyponastic cotyledons of *rhm1* mutants have been proposed to be a consequence of the accumulation of a non-rhamnosylated flavonol (Ringli et al., 2008). As *rhm1-3* alters the flavonol glycosylation profile of flowers, it is possible that the defective petal morphology of *rhm1* mutants is caused by the accumulation of a non-rhamnosylated flavonol. However, most rhamnose in Arabidopsis is located in the cell wall (Reiter et al., 1997), the cell wall is known to control cell morphology, and we have previously shown that *rhm1* mutations decrease the amount of cell wall rhamnose in petals (Saffer et al., 2017). Therefore, we considered the alternative possibility that the *rhm1* petal defects are caused by changes in cell wall rhamnose levels, and that *tt4* suppresses those defects by diverting UDP-L-rhamnose from flavonol glycosylation to the synthesis of cell wall polysaccharides. To test if this hypothesis was plausible, we measured the monosaccharide composition of cell walls from *rhm1-3* and *tt4* single and double mutant flowers. Rhamnose comprised approximately 7.3% of the monosaccharides in cell walls of L*er* wild-type flowers (Figure 3). *rhm1-3* flowers had a substantial decrease in the amount of cell wall rhamnose, with approximately a third less than the wild-type, while the *tt4* mutant was similar to the wild-type. Strikingly, the *rhm1-3 tt4* double mutant flowers were intermediate; there was significantly more cell wall rhamnose than in the *rhm1-3* single mutant, but less than in the wild-type. Therefore, eliminating flavonol biosynthesis redirects UDP-L-rhamnose to the synthesis of cell wall polysaccharides in an *rhm1* mutant background. The only other significant difference in cell wall composition among these mutants was in xylose and mannose levels, which co-eluted in many samples and were combined for these analyses. The percentage of xylose and mannose was increased in *rhm1-3* compared to wild-type or *tt4* (Supplemental Figure 1). This could represent a compensatory change in cell wall composition in *rhm1-3*, or alternatively, when rhamnose levels are decreased then xylose and mannose might comprise a larger fraction of the cell wall without a change in absolute levels.

**Figure 3.**
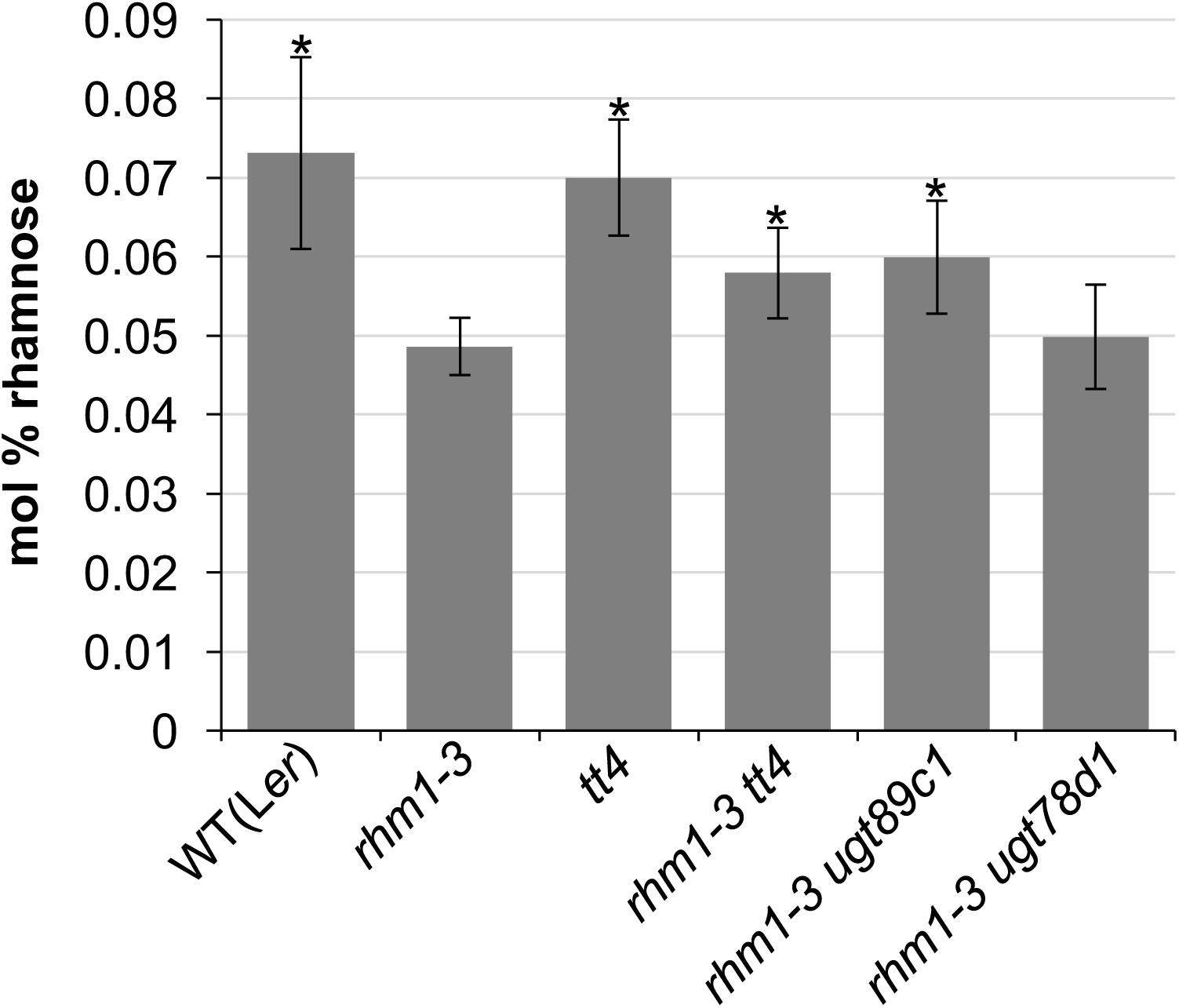
Blocking flavonol synthesis or rhamnosylation increases cell wall rhamnose in an *rhm1* mutant background. Mol % of rhamnose in cell walls from mixed-stage flowers as determined by HPAEC. Error bars show standard deviation and * indicates p<0.05 as compared to *rhm1-3* by t-test. Three independent samples were assayed for each genotype. mol% of other monosaccharides from this experiment are shown in Supplemental Figure 1.

### Blocking flavonol rhamnosylation suppresses the *rhm1* petal defect

Blocking flavonol synthesis could either suppress the *rhm1* petal defects if they are caused by a non-rhamnosylated flavonol, or alternatively if they are caused by a decrease in cell wall rhamnose content that is restored by redirecting rhamnose flux from flavonols to the cell wall. To distinguish between these models, we took advantage of genetic mutants that specifically block flavonol rhamnosylation. *UGT78D1* encodes a flavonol-3-O-rhamnosyltransferase and *UGT89C1* encodes a flavonol-7-O-rhamnosyltransferase (Jones et al., 2003; Yonekura-Sakakibara et al., 2007). *ugt78d1* and *ugt89c1* mutants have a complete loss of flavonol-3-O-rhamnosides and flavonol-7-O-rhamnosides, respectively, indicating that the corresponding genes likely encode the only such enzymes in Arabidopsis (Jones et al., 2003; Kuhn et al., 2016; Yin et al., 2012; Yonekura-Sakakibara et al., 2008; Yonekura-Sakakibara et al., 2007). If the *rhm1* petal defects are caused by accumulation of one or more abnormal non-rhamnosylated flavonols, then blocking flavonol rhamnosylation should cause a similar phenotype. Conversely, since blocking flavonol rhamnosylation prevents rhamnose from being conjugated to flavonols, then we would expect such mutants to redirect rhamnose to the cell wall and rescue the *rhm1* petal defects if they are indeed caused by a lack of cell wall rhamnose.

The *ugt78d1* and *ugt89c1* single mutants had phenotypically normal petals with no helical twisting or hyponasty (Figure 2H-I). The *ugt78d1 ugt89c1* double mutant similarly had normal petals (Figure 2J), despite lacking all flavonol rhamnosyltransferase activity. Together these results indicate that blocking flavonol rhamnosylation does not cause a helical petal phenotype. We next tested if blocking flavonol rhamnosylation could suppress the *rhm1* petal defects. The *rhm1-3 ugt78d1* double mutant had a similar but perhaps somewhat milder twisted petal phenotype as compared to the *rhm1-3* single mutant (Figure 2K). By contrast, *ugt89c1* fully suppressed the *rhm1-3* petal phenotype, as the *rhm1-3 ugt89c1* double mutant had phenotypically normal petals with no helical twisting (Figure 2L). Similar results were obtained in double mutant combinations with *rhm1-2*. The *rhm1-2 ugt78d1* double mutant was indistinguishable from the *rhm1-2* single mutant (Figure 2M). However, *ugt89c1* strongly suppressed the *rhm1-2* hyponastic petal defect, as *rhm1-2 ugt89c1* double mutant petals were nearly wild-type with only slight helical twisting (Figure 2N).

To determine the effects of blocking flavonol rhamnosylation on rhamnose flux, we measured the cell wall monosaccharide composition of *rhm1-3 ugt78d1* and *rhm1-3 ugt89c1* double mutants. The *rhm1-3* single and the *rhm1-3 ugt78d1* double mutant flowers had similar amounts of cell wall rhamnose (Figure 3A). By contrast, the *rhm1-3 ugt89c1* double mutant had less cell wall rhamnose than the wild-type but significantly more than the *rhm1-3* single mutant, similar to the *rhm1-3 tt4* double mutant (Figure 3A). These observations indicate that the *ugt89c1* mutation alters rhamnose flux, partially restoring cell wall rhamnose levels in *rhm1* mutant flowers.

### Blocking flavonol synthesis or rhamnosylation suppresses the *rhm1* mutant cotyledon defect

To determine if the role of flavonols in altering rhamnose flux were tissue-specific or more general, we examined cotyledon morphology in several mutant backgrounds. The cotyledons of *rhm1-2* mutant seedlings have abnormal adaxial pavement cell morphology and consequently are hyponastic (Ringli et al., 2008). Similarly, *rhm1-3* mutant seedlings often have hyponastic cotyledons instead of the epinastic cotyledons seen in wild-type (Figure 4A, C). Correspondingly, *rhm1-3* cotyledons display a cell expansion defect, where the adaxial pavement cells typically fail to form the interlocking lobes characteristic of wild-type pavement cells (Figure 4B, D). *rhm1-2* plants displayed a strongly penetrant hyponastic cotyledon phenotype, while *rhm1-3* plants had a weaker partially penetrant hyponastic cotyledon defect (Table 1). The hyponastic cotyledon phenotype of *rhm1-2* mutants was previously shown to be suppressed by mutations in *tt4* (Ringli et al., 2008). Similarly, we found that *tt4 rhm1-3* double mutants displayed normal cotyledons, indicating that *tt4* completely suppresses the *rhm1-3* hyponastic cotyledon phenotype (Table 1).

**Figure 4.**
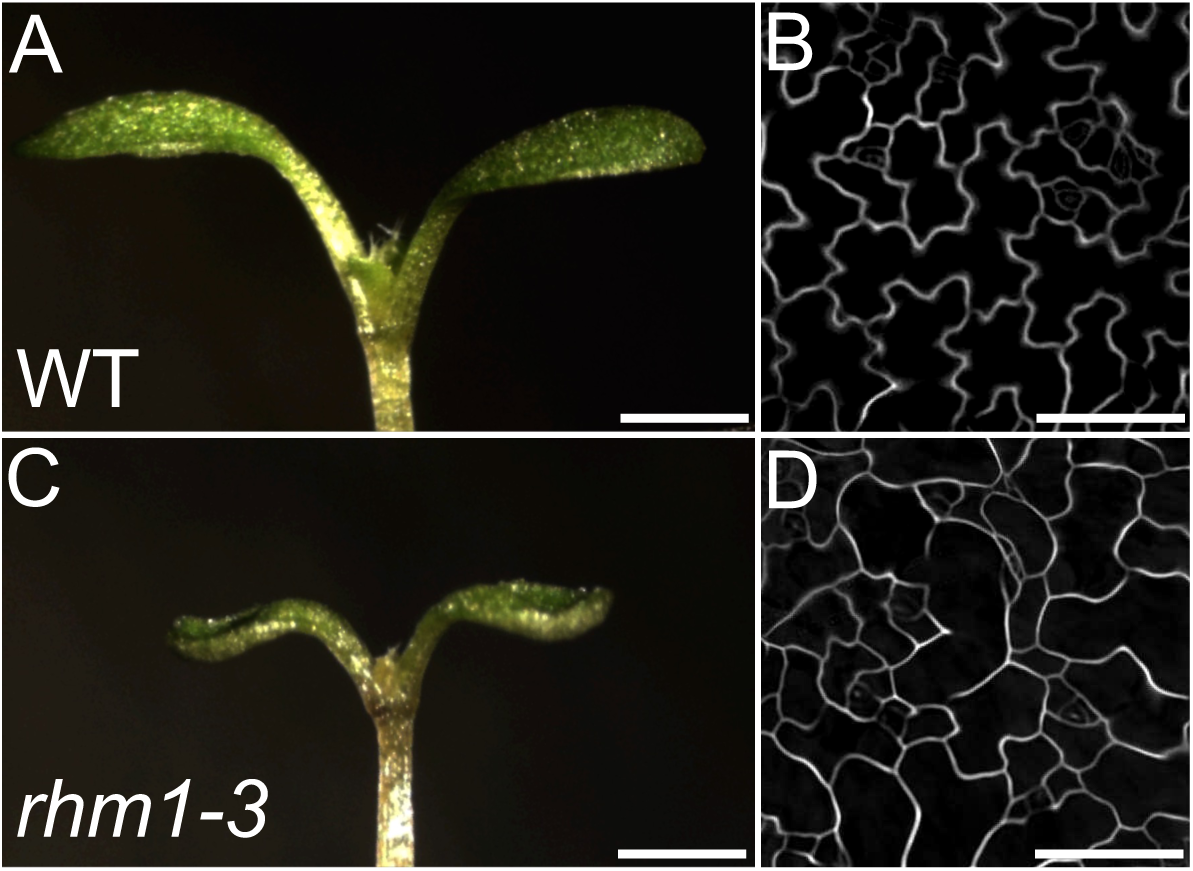
*rhm1-3* cotyledons have defective adaxial cell expansion. All images are from five day old seedlings. Whole seedlings (A and C) and DIC images of agarose impressions of adaxial epidermal cells (B and D) from L*er* wild-type (A and B) and *rhm1-3* (C and D). Scale bars are 1 mm for A, C and 100 μm for B, D.

**Table 1.** The percentage of plants with hyponastic cotyledons was assayed for each genotype on three different days, and the average penetrance ± standard deviation from those three replicates is shown. Seedlings with one hyponastic and one wild-type cotyledon were scored as hyponastic. At least 150 seedlings of each genotype were observed in each replicate. (n) indicates total number of seedlings with either wild-type or hyponastic cotyledons assayed in all three replicates.

| Genotype | % hyponastic cotyledons (n) |
| --- | --- |
| wild-type (Ler) | 1.5 ± 0.6 (672) |
| wild-type (Col) | 0.4 ± 0.4 (862) |
| *rhm1-3* | 47.0 ± 10.1 (929) |
| *rhm1-2* | 82.8 ± 7.5 (982) |
| *tt4* | 1.0 ± 0.8 (550) |
| *rhm1-3 tt4* | 0.1 ± 0.2 (767) |
| *ugt89c1* | 0.2 ± 0.4 (922) |
| *ugt78d1* | 1.0 ± 0.2 (780) |
| *ugt89c1 ugt78d1* | 0.6 ± 0.5 (880) |
| *rhm1-3 ugt89c1* | 0.4 ± 0.4 (789) |
| *rhm1-3 ugt78d1* | 22.4 ± 6.0 (915) |
| *rhm1-2 ugt89c1* | 1.2 ± 1.0 (863) |
| *rhm1-2 ugt78d1* | 76.6 ± 6.0 (712) |

The hyponastic cotyledons seen in *rhm1-3* and *rhm1-2* mutants were previously proposed to be due to the presence of a non-rhamnosylated flavonol (Ringli et al., 2008). Alternatively, as is the case in petals, the hyponastic cotyledon phenotype in *rhm1* mutants might be due to a decrease in the amount of rhamnose in the cell wall, with *tt4* suppressing that defect by redirecting rhamnose from flavonols to the cell wall. We distinguished between these alternative hypotheses by genetically blocking flavonol rhamnosylation.

Neither the *ugt78d1* or *ugt89c1* mutations resulted in a hyponastic cotyledon phenotype, indicating that blocking either 3-O-or 7-O-rhamnosylation of flavonols does not cause an *rhm1*-like phenotype in seedlings (Table 1). Furthermore, the *ugt78d1 ugt89c1* double mutant also displayed wild-type cotyledons, suggesting that accumulation of non-rhamnosylated flavonols does not cause hyponastic growth of cotyledons. Instead, *ugt89c1* fully suppressed the hyponastic cotyledon phenotype of both *rhm1-3* and *rhm1-2*. *ugt78d1* partially suppressed the penetrance of hyponastic cotyledons in *rhm1-3*, but did not appreciably alter the penetrance of hyponastic cotyledons in the more severe *rhm1-2* mutant. These results are in agreement with previously reported single and double mutant phenotypes (Kuhn et al., 2016; Ringli et al., 2008). Together, these observations indicate that blocking flavonol-3-O-rhamnosylation only slightly suppressed the *rhm1* cotyledon phenotype, while blocking flavonol-7-O-rhamnosylation fully suppressed the cotyledon defect.

To determine if the different degree of suppression by *ugt78d1* and *ugt89c1* correlated with the amount of rhamnose at the 3 and 7 positions of flavonols that could be redirected to other molecules, we re-analyzed the previously reported flavonol glycosylation profiles of wildtype and *rhm1-2* mutant seedlings. Ringli *et al.* (2008) reported the area of each flavonol glycoside in shoots from wildtype and *rhm1* mutant seedlings, and we summed the peak areas of individual flavonol glycosides to determine the percentage of flavonols that were 3-O-and 7-O-rhamnosylated (Supplemental Table 1). Nearly all identified flavonol glycosides from wild-type seedlings were 7-O-rhamnosylated, while approximately two thirds were 3-O-rhamnosylated. Similar to the effect of *rhm1-3* in petals, *rhm1-2* had a more severe effect on 3-O-rhamnosylation than on 7-O-rhamnosylation in seedlings, such that while there was a decrease in both 3-O-and 7-O-rhamnosides, the ratio of 7-O-rhamnosides to 3-O-rhamnosides in *rhm1-2* seedlings was increased to 2.75:1. Therefore in *rhm1-2* seedlings there is substantially more flavonol-7-O conjugated rhamnose than flavonol-3-O conjugated rhamnose.

### *rhm1* mutant seedlings have decreased cell wall rhamnose levels

To determine if *rhm1* mutants affect the cell wall of seedlings, we assayed the monosaccharide composition of cell walls isolated from the aerial portion of seedlings from *rhm1-3*, *rhm1-1* (also known as *rol1-1*), and *rhm1-2* mutants and the corresponding wild-types (Figure 5). Each *rhm1* mutant had a significant decrease in the amount of cell wall rhamnose compared to the wild-type, ranging from a 10% decrease in *rhm1-1* to a 23% decrease in *rhm1-3*. Several other monosaccharides differed between the wild-type and specific *rhm1* mutant alleles. For example, both *rhm1-1* and *rhm1-2* had an increased percentage of xylose as compared to wild type. However, rhamnose was the only monosaccharide that was significantly affected in all three *rhm1* alleles as compared to the wild-type.

**Figure 5.**
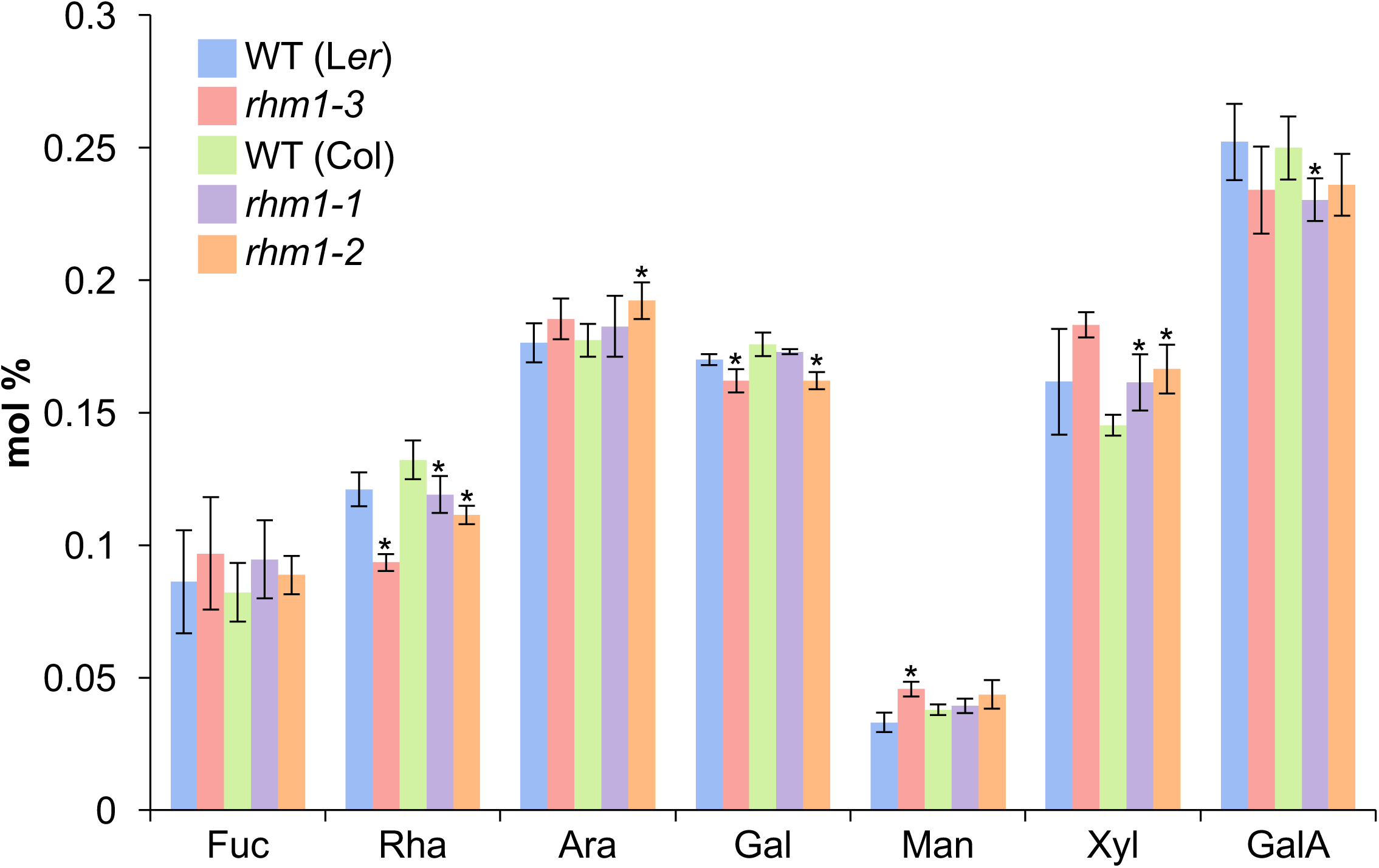
*rhm1* mutant seedlings have decreased cell wall rhamnose content. Monosaccharide composition of cell walls isolated from the aerial tissues of five day old seedlings. Fucose (Fuc), Rhamnose (Rha), Arabinose (Ara), Galactose (Gal), Xylose (Xyl), Mannose (Man), Galacturonic acid (GalA). Four independent samples were assayed for each genotype. Error bars show standard deviation and * indicates p<0.05 as compared to the corresponding wild-type by t-test.

### The *rhm1* cotyledon defect is enhanced by mutants with altered pectin

Because rhamnose is primarily located in the pectic fraction of cell walls, we investigated cotyledon development in mutants that alter pectin structure or abundance. RG-II is a complex pectic polysaccharide containing at least 12 distinct sugar residues including both rhamnose and fucose. *MURUS1* (*MUR1)* encodes a GDP-D-mannose-4,6-dehydratase that is required for synthesis of L-fucose and *mur1* mutants lack nearly all fucose in aerial organs, causing an abnormal RG-II structure that impairs its borate-mediated crosslinking (Bonin et al., 1997; O’Neill et al., 2001; Reiter et al., 1993; Reuhs et al., 2004). *QUASIMODO1 (QUA1)* encodes a glycosyltransferase and *qua1* mutants have decreased homogalacturonan levels (Bouton et al., 2002). The *mur1* mutant caused a weak hyponastic cotyledon defect with 5% penetrance (Table 2). Most *qua1* seedlings had severe cell adhesion defects that precluded categorizing them as either wild-type or hyponastic, but the assayable *qua1* seedlings with less severe cell adhesion defects did not display hyponastic cotyledons (Table 2). To test if RG-II and HG have a subtle role in cotyledon development, we next tested if *mur1* and *qua1* could modify the partially penetrant hyponastic cotyledon phenotypes of *rhm1-3* and *rhm1-1* seedlings. We observed that 84% of *rhm1-3* seedlings had hyponastic cotyledons, while nearly all *rhm1-3 mur1* double mutant cotyledons were hyponastic. Similarly, only 6% of *rhm1-1* seedlings displayed hyponastic cotyledons, while nearly half of *rhm1-1 mur1* double mutant seedlings had hyponastic cotyledons. *rhm1 qua1* double mutants had less severe cell adhesion defects than the *qua1* single mutant; it is unclear if *rhm1* can suppress the *qua1* cell adhesion defect or if the reduced severity is a consequence of the *rhm1* mutants being in a different ecotype background than *qua1*. The *qua1* mutation strongly enhanced the hyponastic cotyledon defect of *rhm1* mutants, with 96% of *rhm1-3 qua1* double mutants and 89% of *rhm1-1 qua1* double mutant seedlings displaying hyponastic cotyledons. The enhancement of the *rhm1* cotyledon phenotype by *mur1* and *qua1* indicates that alterations in pectin composition can promote hyponastic growth of cotyledons.

**Table 2.** The percentage of plants with hyponastic cotyledons was assayed for each genotype on three different days, and the average penetrance ± standard deviation from those three replicates is shown. Seedlings with one hyponastic and one wild-type cotyledon were scored as hyponastic. Many plants carrying the *qua1* mutation exhibited severe cell adhesion defect that precluded that assaying of hyponastic vs wild-type cotyledon morphology. At least 100 seedlings of each genotype were observed in each replicate, except for the *qua1* single mutant. (n) indicates total number of seedlings with either wild-type or hyponastic cotyledons assayed in all three replicates.

| Genotype | % hyponastic cotyledons (n) |
| --- | --- |
| wild-type (Ler) | 1.5 ± 0.4 (1220) |
| wild-type (Col) | 0.2 ± 0.3 (1067) |
| wild-type (Ws) | 0.1 ± 0.2 (1021) |
| *rhm1-3* | 83.8 ± 4.9 (1297) |
| *rhm1-1* | 6.0 ± 2.2 (1096) |
| *mur1* | 4.7 ± 1.5 (1224) |
| *qua1* | 0.8 ± 1.4 (104) |
| *rhm1-3 mur1* | 99.0 ± 0.6 (1249) |
| *rhm1-1 mur1* | 46.7 ± 8.4 (1192) |
| *rhm1-3 qua1* | 95.7 ± 1.8 (786) |
| *rhm1-1 qua1* | 89.3 ± 2.1 (457) |

## Discussion

### Alternative models for the effect of flavonols on *rhm1* mutant phenotypes

In plant cells, the primary classes of molecules known to contain rhamnose are flavonols and the cell wall polysaccharides RG-I and RG-II. We and others have shown that *rhm1* mutations decrease both flavonol rhamnosylation and cell wall rhamnose in seedlings and flowers (Figure 1; Figure 5; (Saffer et al., 2017)(Ringli et al., 2008; Yonekura-Sakakibara et al., 2008)). *rhm1* mutants have abnormal epidermal cell morphology, including cotyledon pavement cells, conical petal epidermal cells, trichomes, and root hairs (Diet et al., 2006; Ringli et al., 2008; Saffer et al., 2017). Each of these developmental defects could be caused by decreased cell wall rhamnose or flavonol rhamnosylation. Based on genetic and biochemical analyses, we have previously proposed that the cell expansion and helical growth defects in *rhm1* petal cells are a consequence of decreased levels of a rhamnose-containing cell wall polymer, most likely RG-I (Saffer et al., 2017). In this study, we show that decreased levels of cell wall rhamnose cause the *rhm1* cell morphology defects, and blocking flavonol rhamnosylation can suppress those defects by diverting rhamnose from flavonols to the cell wall. Conversely, it has been proposed that the *rhm1* phenotype in cotyledons is due to accumulation of non-rhamnosylated flavonols (Kuhn et al., 2016; Kuhn et al., 2011; Ringli et al., 2008). However, there are a variety of observations that are better explained by a model in which the *rhm1* epidermal cell morphology defects are caused by decreased cell wall rhamnose than by the accumulation of non-rhamnosylated flavonols.

It has been proposed that decreased flavonol rhamnosylation in *rhm1* mutants causes morphological defects by altering auxin signaling (Kuhn et al., 2016; Kuhn et al., 2011; Ringli et al., 2008), but the evidence cited in support of this hypothesis leaves open the possibility that changes in auxin signaling might not cause the *rhm1* cotyledon defects. This model was supported by the observation that wild type seedlings treated with the auxin transport inhibitor 1-N-Naphthylphthalamic acid (NPA) have hyponastic cotyledons similar to *rhm1* mutants (Ringli et al., 2008). However, unlike *rhm1* mutants, NPA-treated seedlings displayed wild-type adaxial pavement cell morphology (Ringli et al., 2008). Mutations affecting many other processes, including adaxial-abaxial polarity, maintenance of cell divisions, epidermal cell differentiation, and miRNA biogenesis can cause hyponastic growth of leaves and cotyledons (Ha et al., 2007; Han et al., 2004; Lu and Fedoroff, 2000; Qin et al., 2005; Tao et al., 2013; Wu et al., 2005), and so the failure of NPA to fully phenocopy the cotyledon morphology of *rhm1* suggests that NPA and *rhm1* cause hyponastic cotyledon phenotypes by distinct mechanisms.

The *rhm1* hyponastic cotyledon phenotype is suppressed by *ugt89c1* mutants that blocks flavonol-7-O-rhamnosylation and by *fls1* mutants that block flavonol synthesis (Kuhn et al., 2016; Kuhn et al., 2011)(Table 1). To explain the suppression by both *ugt89c1* and *fls1*, two distinct models were proposed to integrate conflicting results. In a protoplast assay, *rhm1* increased transport of the auxin naphthalene-1-acetic acid (NAA), and incorporation of an *fls1* mutation restored wild-type NAA transport levels (Kuhn et al., 2011), suggesting that altered auxin transport could be responsible for the *rhm1* cotyledon defects. However, *ugt89c1* suppresses the abnormal cotyledon morphology of *rhm1*, but not the *rhm1* auxin transport defect (Kuhn et al., 2016).

Instead, *ugt89c1* alters the levels of several auxin precursors and degradation intermediates, leading to the hypothesis that the effects of *ugt89c1* on the levels of these molecules explains the *ugt89c1* suppression of *rhm1* cotyledon defects independent of auxin transport (Kuhn et al., 2016). For three out of five auxin-related compounds tested, *ugt89c1* and *fls1* had opposite effects on their abundance, despite both *ugt89c1* and *fls1* having equivalent phenotypic effects on *rhm1* cotyledon development. *ugt89c1* and *fls1* mutants both decrease the level of rhamnosylated flavonols and suppress *rhm1* cell morphology defects, but have different effects on auxin signaling, raising the possibility that their roles in auxin signaling might not be related to their suppression of the *rhm1* phenotype.

### The degree of *rhm1* suppression corresponds to the amount of rhamnose diverted from flavonols

The amount of rhamnose that *tt4*, *ugt89c1*, and *ugt78d1* can potentially divert from flavonols to the cell wall correlates with how well each mutation suppresses *rhm1* epidermal cell defects. The *tt4* mutation, which fully suppresses *rhm1* cotyledon and petal defects, prevents flavonol synthesis and consequently all flavonol-conjugated rhamnose can be diverted to the cell wall. A larger percentage of flavonols are 7-O-rhamnosylated than are 3-O-rhamnosylated, and *rhm1* mutants cause more severe reductions in the abundance of flavonol-3-O-rhamnosides than flavonol-7-O-rhamnosides (Supplemental Table 1)(Yonekura-Sakakibara et al., 2008). The *ugt89c1* mutation can therefore block most but not all flavonol rhamnosylation in *rhm1* mutants. Accordingly, *ugt89c1* fully suppresses the petal and cotyledon defects of *rhm1-3* and the cotyledon defect of the stronger *rhm1-2* allele, and can largely suppress the *rhm1-2* petal phenotype. Only a small portion of flavonol rhamnose in *rhm1* mutants is 3-O-conjugated, and so *ugt78d1* has little effect on most *rhm1* mutant phenotypes, but can partially suppress the *rhm1-3* cotyledon phenotype. Consistent with the amount of flavonol-conjugated rhamnose that can be diverted, blocking flavonol-7-O-rhamnosylation in a *rhm1-3* background redirects nearly as much rhamnose to the cell wall as blocking all flavonol synthesis, while blocking flavonol-3-O-rhamnosylation has no obvious effect on cell wall rhamnose levels (Figure 3).

### Flavonols modify *rhm1* defects by altering rhamnose flux

The suppression of *rhm1* epidermal cell defects by eliminating flavonols is consistent either with the defects resulting from the accumulation of a non-rhamnosylated flavonol, or with a model where blocking flavonol synthesis diverts rhamnose to a different rhamnose-containing macromolecule required for the morphogenesis of epidermal cells. However, mutants that fully block either flavonol-3-O or flavonol-7-O rhamnosylation do not develop an *rhm1*-like phenotype. Rather, blocking flavonol rhamnosylation suppresses the helical petals, abnormal pavement cell morphology, and hyponastic cotyledons of *rhm1* (Figure 2; Table 1)(Kuhn et al., 2016). Both *rhm1* and *ugt89c1* mutations decrease the abundance of rhamnosylated flavonols, yet have antagonistic effects on epidermal cell morphology, which is inconsistent with the proposition that the *rhm1* phenotype results from the accumulation of a non-rhamnosylated flavonol.

Instead, the suppression of *rhm1* defects by both *tt4* and *ugt89c1* is consistent with a model in which blocking flavonol synthesis or rhamnosylation diverts rhamnose from flavonols to other rhamnose-containing molecules to restore normal development to *rhm1* mutants. This hypothesis is further supported by measurements of cell wall rhamnose levels, which indicate that preventing flavonol synthesis or rhamnosylation in an *rhm1* mutant background redirects UDP-L-rhamnose to the synthesis of cell wall polysaccharides (Figure 3). Thus, we propose that flavonol glycosylation indirectly affects epidermal cell development in *rhm1* mutants by altering rhamnose flux (Figure 6). In *rhm1* mutants there is a decreased pool of UDP-L-rhamnose available for flavonol glycosylation and synthesis of cell wall polysaccharides (Figure 6B). By this model, the decreased levels of rhamnose-containing cell wall polysaccharides cause the abnormal epidermal cell morphology seen in *rhm1* petals and cotyledons. Blocking flavonol synthesis (with *tt4* or *fls1*) or rhamnosylation (with *ugt89c1*) prevents rhamnose from being conjugated to flavonols, allowing for rhamnose to be diverted to the cell wall, which restores sufficient levels of cell wall rhamnose to permit normal development (Figure 6C). In plants with compromised UDP-L-rhamnose synthesis, flavonols can affect development by altering rhamnose flux, but altered flavonol rhamnosylation is not directly responsible for the abnormal morphology of *rhm1* mutant petal or cotyledon epidermal cells. This model provides a single consistent explanation for the suppression of *rhm1* mutant phenotypes by *tt4*, *fls1*, and *ugt89c1* mutants.

**Figure 6.**
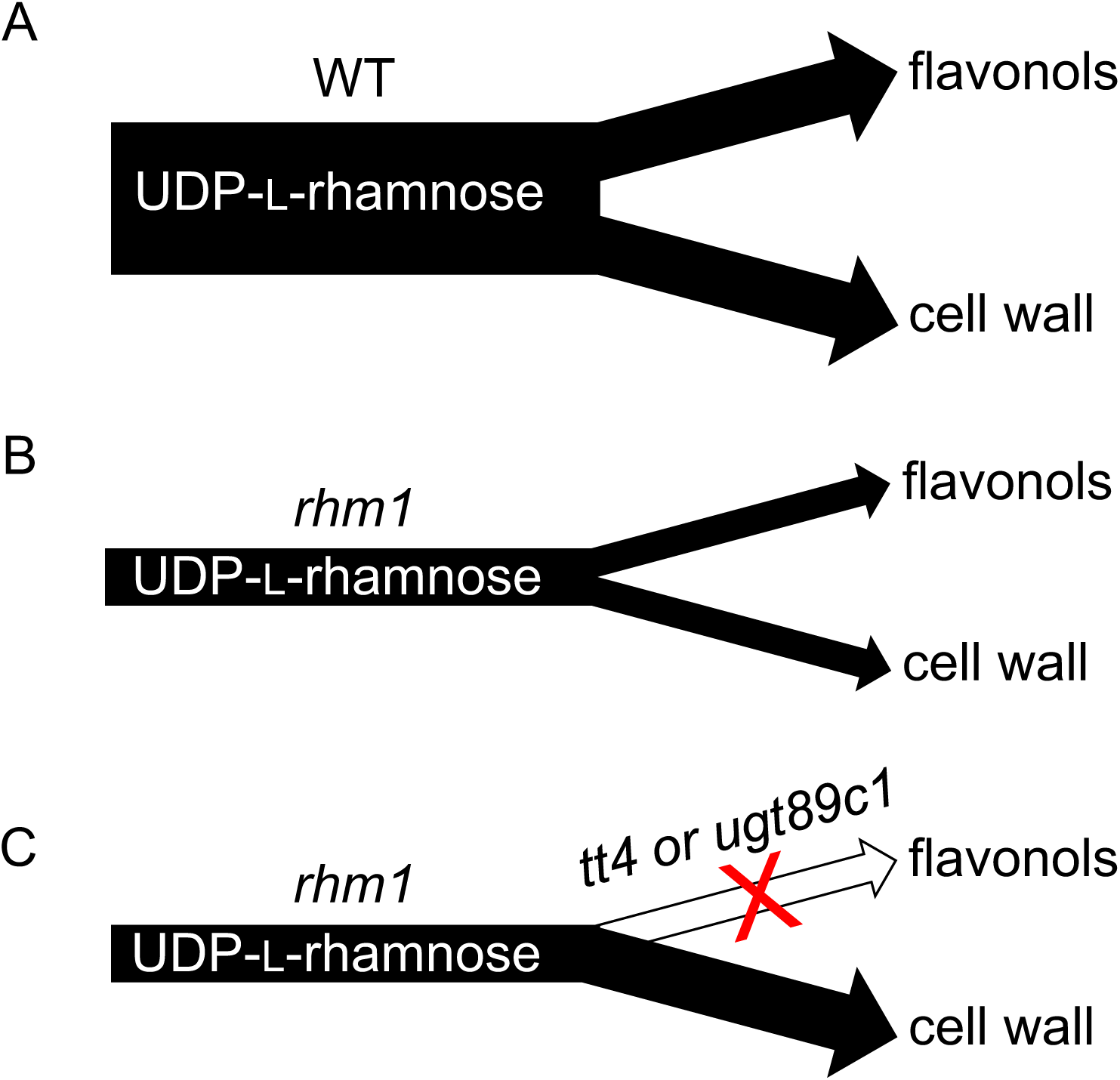
Model-Flavonols affect *rhm1* phenotype by altering rhamnose flux. A) In the wild-type, UDP-L-rhamnose is primarily directed to the synthesis of the cell wall polysaccharides RG-I and RG-II and the glycosylation of flavonols. B) In *rhm1* mutants, the loss of a UDP-L-rhamnose synthase decreases the pool of UDP-L-rhamnose, reducing the synthesis of cell wall polysaccharides and the rhamnosylation of flavonols. C) Blocking either the synthesis or rhamnosylation of flavonols (with *tt4* or *ugt89c1* mutants, respectively) diverts the rhamnose that would normally have been conjugated to flavonols to the cell wall, partially or fully restoring normal cell morphology to *rhm1* mutants.

The effects of flavonol synthesis and rhamnosylation on rhamnose flux in *rhm1* raises the intriguing possibility that flavonols might normally function as a natural buffering pool of rhamnose. Our results indicate that cell wall rhamnose is essential for proper development, while mutants lacking flavonols are viable (Burbulis et al., 1996) and some rhamnosylated flavonols might only be important under specific conditions such as for herbivore defense (Onkokesung et al., 2014). Perhaps the presence of a pool of rhamnose directed to non-essential flavonols can act as a buffer to prevent fluctuations in UDP-L-rhamnose synthesis or in the rate of cell wall expansion from causing deleterious changes to cell wall rhamnose levels.

### Pectin promotes normal cotyledon development

Cell expansion in plants is driven by turgor pressure exerting force on the entire cell, and so the shape of plant cells is controlled by their cell wall (Cosgrove, 2005). *rhm1* mutants decrease cell wall rhamnose levels in multiple tissues (Figure 5; (Saffer et al., 2017)), likely by affecting the synthesis of the pectic polysaccharides RG-I and RG-II (Diet et al., 2006; Saffer et al., 2017). The *mur1* mutant has defective RG-II structure (O’Neill et al., 2001) and occasionally exhibits hyponastic cotyledons (Table 2), but at a much lower penetrance than strong *rhm1* alleles, indicating that while RG-II does prevent hyponastic growth of cotyledons, the *rhm1* cotyledon defects are probably not primarily caused by an RG-II defect. Therefore, the hyponastic cotyledons of *rhm1* mutants are likely primarily a consequence of decreased levels of RG-I, although changes in RG-II levels or structure might also contribute to the phenotype. The *qua1* mutant is partially defective in HG synthesis (Bouton et al., 2002), but does not cause hyponastic cotyledons, suggesting that HG is less important than RG-I or RG-II for preventing hyponastic cotyledon growth. Both *mur1* and *qua1* can enhance the *rhm1* hyponastic cotyledon defect, consistent with *rhm1* hyponasty resulting from altered pectin composition. Together, these results indicate that multiple pectic polysaccharides contribute to differing degrees to the morphogenesis of cotyledon epidermal cells, with RG-I likely having a major role in promoting cell expansion and preventing hyponasty.

### Cell wall rhamnose is broadly required for epidermal cell morphology

Blocking flavonol synthesis suppresses the effect of *rhm1* mutants on the morphology of cotyledons and petals but not root hairs (Ringli et al., 2008). Given the effects of flavonols on rhamnose flux, we suggest that *RHM1* has a comparable role in controlling cell morphology through maintaining the levels of cell wall rhamnose in petals, cotyledons, and roots, and that the cell-type specific suppression of *rhm1* defects by *tt4* could derive from different ratios of rhamnose directed to flavonols versus pectic polysaccharides in each tissue. *RHM1* is one of three highly similar UDP-L-rhamnose synthases in Arabidopsis (Reiter and Vanzin, 2001), and the specific tissues affected by *rhm1* mutants might reflect tissue-specific requirements for *RHM1*, given the different expression patterns of the *RHM* genes and tissue-specific requirements for flavonol rhamnosylation. Consistent with this, *RHM1*, *UGT89C1*, and *FLS1* are all predominantly expressed in adaxial epidermal cells where *rhm1* phenotypic abnormalities are observed (Kuhn et al., 2016; Kuhn et al., 2011; Saffer et al., 2017). Thus, we propose that the *rhm1* phenotype reflects a broadly important role for rhamnose-containing pectic polysaccharides in the morphogenesis of epidermal cells.

## Materials and Methods

### Plant growth

Plants were grown in a mix of two parts vermiculite to one part Fafard Superfine Germinating Mix soil for assays of petals and on solid media consisting of half-strength Murashige and Skoog (MS) salts, 2% (w/v) sucrose, 100 mg/L myo-inositol, and 0.6% (w/v) phytagel (Sigma) for assays of cotyledon development. All plants were grown under long day conditions (16 hr light/8 hr dark) at 22°C. Images were obtained with an AxioCam HRc connected to a Zeiss Stemi 2000-C dissecting microscope.

### Genetic material

Three alleles of *RHM1* are used in this study, the nonsense mutant *rhm1-1* (also known as *rol1-1*), the missense mutant *rhm1-2* (also known as *rol1-2*), and the missense mutant *rhm1-3* (Diet et al., 2006; Saffer et al., 2017). For simplicity, in this manuscript we refer to the previously identified *rhm1* alleles *rol1-1* and *rol1-2* as *rhm1-1* and *rhm1-2,* respectively. The *tt4* mutant used in this study is *tt4-1* (also known as W85), the *ugt89c1* mutant is the T-DNA insertion SALK_068559, and the *ugt78d1* mutant is the T-DNA insertion SAIL_568_F08 (Alonso et al., 2003; Jones et al., 2003; Sessions et al., 2002; Shirley et al., 1995; Yonekura-Sakakibara et al., 2007). To perturb cell wall composition the *qua1-1* and *mur1-1* mutations were used (Bouton et al., 2002; Reiter et al., 1993). *rhm1-3* and *tt4-1* are derived from the L*er* ecotype, *qua1-1* is derived from the Ws ecotype, and all other mutants in this study are derived from the Col ecotype. *qua1-1* was a gift from Grégory Mouille, the isolation of the *rhm1-3* mutation is described (Saffer et al., 2017), and all other seed stocks were obtained from the Arabidopsis Biological Resource Center.

### Genotyping and double mutant construction

Genotyping PCR assays for *rhm1-3*, *mur1-1*, and *qua1-1* (Saffer et al., 2017), and *rhm1-1* and *rhm1-2* (Diet et al., 2006) were previously described. *tt4-1* abolishes a BsaI restriction site, and the presence of the *tt4-1* mutation was detected by amplifying DNA with the primers 5’-TCGGTCAGGCTCTTTTCAGT-3’ and 5’-TGTCGCCCTCATCTTCTCTT-3’ and digesting with BsaI. Homozygous *tt4-1* plants were confirmed by their transparent testa phenotype. For the SAIL_568_F08 T-DNA insertion, the primer 5’-GACATGCATGCTAACAGATGC-3’ was used in combination with either 5’-GCTTCCTTTCATGGAGAAATC-3’ to detect wild-type DNA or 5’-GCCTTTTCAGAAATGGATAAATAGCCTTGCTTCC-3’ to detect the T-DNA insertion. For the SALK_068559 T-DNA insertion, the primer 5’-ACACGTTTCCTGAATCACCAC-3’ was used in combination with either 5’-ACCGCGTGTGTAATGTATCG-3’ to detect wild-type DNA or 5’-ATTTTGCCGATTTCGGAAC-3’ to detect the T-DNA insertion.

### Cotyledon epidermal cell morphology

Seedlings were placed adaxial side down in a molten solution of 3% (w/v) low melting point agarose in water. The agarose was allowed to cool until solid and the seedling was removed. The cotyledon impression in the agarose was visualized using DIC optics on a Zeiss Axiophot microscope at 10x magnification.

### Identification and quantification of flavonol glycosides

Each sample consisted of approximately 150 mg fresh weight of mixed stage flowers, which was equivalent to approximately 20 mg dry weight (DW). Three independent samples were assayed for each genotype. Tissue was frozen in liquid N2, lyophilized overnight, and then homogenized using a Qiagen TissueLyser and 5 mm steel balls. 1μL of a 1 mM solution of the isoflavone daidzein per mg of DW was added to each sample as an internal standard. 20 μL methanol per mg DW was added to each sample, which were then sonicated for one hour at room temperature to extract methanol-soluble compounds. After centrifugation the methanol extract was transferred to a new tube, dried under vacuum, and resuspended in 20 μL 50% aqueous methanol per mg DW, and then particulates were removed with a 0.45μm filter (Millipore MultiScreen MSHVN4510). HPLC was performed on a Dionex UltiMate 3000 HPLC with a C18 column (Ace 111-1503, 150 mm length and 3.0 mm inner diameter with 3 μm particle size) with a flow rate of 0.5 mL/min at 30°C. 10 μL of each sample was injected.

Compounds were separated by elution with gradients of solvent A (water with 0.1% formic acid) and solvent B (90% acetonitrile and 10% water with 0.1% formic acid). The column was first equilibrated with 90% solvent A and 10% solvent B for 5 min. Then starting from 90% solvent A and 10% solvent B at time 0, the solvent concentration was changed in linear gradients to 80% solvent A and 20% solvent B at 15 min, 60% solvent A and 40% solvent B at 25 min, 100% solvent B at 30 min, and then held at 100% solvent B until 35 min. UV absorbance was assayed with a DAD3000 diode array detector with detectors set to 210 nm, 254 nm, 280 nm, and 320 nm. Flavonol abundances were determined from absorbance at 280 nm by calculating the area under each peak using Chromeleon software version 6.8 (Dionex). The HPLC was connected to a MSQ Plus mass spectrometer set to use electrospray ionization (ESI) in both positive-and negative-ionization modes with a 500°C probe temperature, 50V cone voltage, and mass range of 100-1000. Peaks likely to represent flavonols were identified based on their characteristic UV absorbance spectra with maxima at approximately 260 nm and 370 nm (de Rijke et al., 2006). Some peaks represented different compounds in *rhm1-3 tt4* samples compared to either L*er* or *rhm1-3* samples; a given flavonol glycoside was considered not detected in a sample if the peak had a non-flavonol-like UV absorbance spectra and lacked the appropriate mass ion. Peaks were assigned to specific flavonol glycosides on the basis of mass and comparison of mass and order of retention to a previously published analysis of flavonol glycosides in Arabidopsis flowers (Yonekura-Sakakibara et al., 2008).

### Monosaccharide composition of cell walls

Monosaccharide composition of alcohol insoluble residue (AIR) was determined by high performance anion exchange chromatography with pulsed amperometric detection (HPAEC-PAD) essentially as described (Saffer et al., 2017). For analysis of cell wall composition of flowers approximately 35 mg fresh weight of mixed stage flowers were collected for each sample, which produced approximately 3 mg AIR. For analysis of cell wall composition of seedlings, seedlings were cut in half and only the aerial tissues were collected. Each seedling sample was derived from approximately 30 mg fresh weight of tissue, which produced approximately 2 mg AIR. Neutral sugars were eluted with either 8 mm NaOH (Figure 3) or 14 mm NaOH (Figure 5). Molar amounts of fucose, rhamnose, arabinose, galactose, mannose, xylose, and galacturonic acid were determined and used for mol% calculations. In the analysis of flower cell wall composition mannose and xylose often co-eluted and were combined.

## Acknowledgments

Grégory Mouille and the Arabidopsis Biological Resource Center kindly provided seed stocks. We thank William Chezem, Teresa Ceserani, and Nicole Clay for assistance with chromatography. A.M.S. was supported in part through a Yale University Brown Fellowship. This project was supported by NSF grant MCB-1615387 to V.F.I.

